# Gatekeeper transcription factors regulate switch between lineage preservation and cell plasticity

**DOI:** 10.1101/2020.03.19.999433

**Authors:** Tania Ray, Anit Shah, Gary A. Bulla, Partha S. Ray

## Abstract

Reprogramming somatic cells to pluripotency by repressing lineage-instructive transcription factors (TFs) alone has not been pursued because lineage specification is thought to be regulated by transcriptional regulatory networks (TRNs) comprising of multiple TFs rather than by single pivotal “gatekeeper” TFs. Utilizing an intra-species somatic cell hybrid model, we identified Snai2 and Prrx1 as the most critical determinants of mesenchymal commitment in rat embryonic fibroblasts (REFs) and demonstrate that siRNA-mediated knockdown of either of these master regulators is adequate to convert REFs into functional adipocytes, chondrocytes or osteocytes without requiring exogenous TFs or small molecule cocktails. Furthermore, knockdown of Snai2 alone proved sufficient to transform REFs to dedifferentiated pluripotent stem-like cells (dPSCs) that formed embryoid bodies capable of triple germ-layer differentiation. These findings suggest that inhibition of a single gatekeeper TF in a lineage committed cell is adequate for acquisition of cell plasticity and reprogramming without requiring permanent genetic modification.

**Graphical Abstract:** 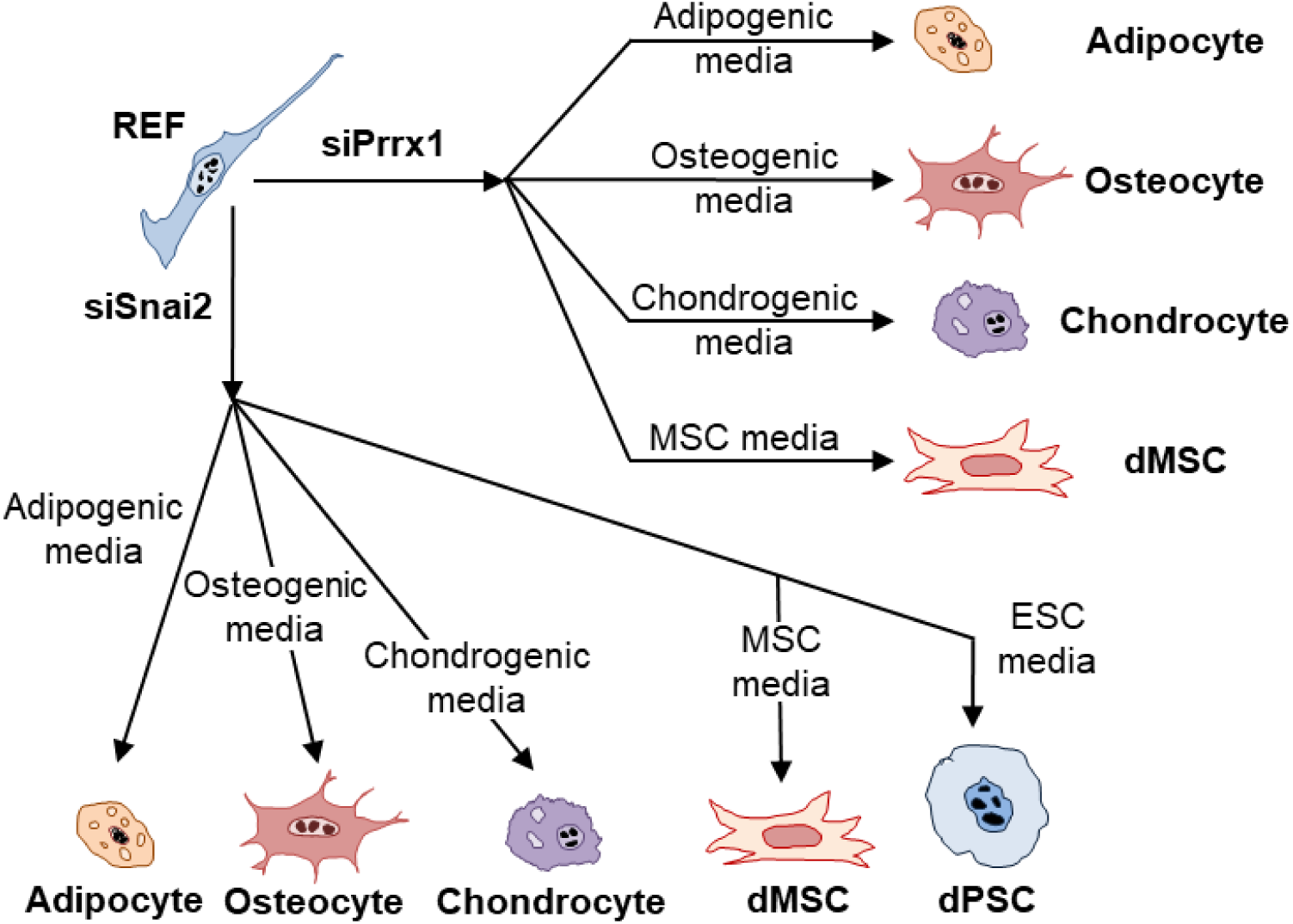

Schematic diagram depicting transdifferentiation of REFs into adipocytes, osteocytes, chondrocytes and dedifferentiation into MSCs on individual treatment with siSnai2 or siPrrx1. dPSCs were generated only in the siSnai2 group.

## Introduction

Direct reprogramming of cells to pluripotency or alternate lineages from easily available somatic cells has proven the fact that cell differentiation is not a unidirectional and irreversible process. In 2006, researchers Takahashi and Yamanaka revolutionized stem cell research by showing that forced expression of only four exogenous Transcription Factors (TFs) Oct4, Sox2, Klf4, and c-Myc (OSKM) was sufficient to convert fibroblast cells into embryonic stem cell-like cells, which they named induced pluripotent stem cells (iPSCs). Since this ground breaking discovery (Takahashi and Yamanaka, 2006), several studies have further expanded this work (Graf, 2011; Srivastava and DeWitt, 2016). Many groups have also reported direct programming of lineage-committed somatic cells into alternate differentiated cell types by forced expression of lineage-instructive TFs (Davis et al., 1987; Vierbuchen et al., 2010; Xu et al., 2015). More recently, small molecule driven chemical reprogramming of somatic cells has been demonstrated to be another strategy to transform differentiated cells (Cao et al., 2016; Liu et al., 2016; Thoma et al., 2014; Xie et al., 2017).

The above reported methods for reprogramming differentiated cells into iPSCs or other cell types often lack efficiency which is thought to be primarily due to retention of the starting cell’s lineage-instructive transcriptional regulatory network (TRN) enforced by lineage-instructive TFs (Hikichi et al., 2013) and the underlying epigenetic state (Nashun et al., 2015). Although significant advances have been made in cell reprogramming strategy, there are no prior reports that demonstrate reprogramming of adult cells into induced pluripotent stem cells (iPSCs) or alternate differentiated cell types by transiently repressing a single pivotal master regulatory lineage-preserving TF of the starting cell alone, in the absence of exogenous TFs (e.g. OSKM) or small molecule cocktails. This is because lineage commitment is known to be regulated by the starting cell’s TRN comprising of multiple TFs working in conjunction to exert transcriptomic and epigenetic control, to enforce and maintain the cell lineage. Despite several clues to the contrary in the scientific literature, the existence of single pivotal “gatekeeper” TFs (GTFs) within the TRN of a lineage-committed cell which could single-handedly regulate the switch between lineage preservation and cell plasticity has never been demonstrated. As such the concept that a critically important biologic function like enforcement of lineage commitment can be disrupted by manipulating a single key TF member of the starting cell’s TRN was not thought to be possible.

Lineage-instructive TFs of differentiated cells strictly enforce cell identity, resist plasticity acquisition and contribute to low reprogramming efficiency associated with exogenous TF-induced and/or small molecule-induced cellular reprogramming. We reasoned that if presence of residual epigenetic memory enforced by master regulatory lineage-instructive TFs of the starting cell is truly a critical impediment to acquisition of cellular plasticity, then lifting these restrictions imposed by such TFs should allow cellular plasticity to be acquired and thereby permit cell reprogramming to proceed unhindered, even in the absence of Yamanaka factors or pathway modulatory cocktails. We further hypothesized that among a cell’s lineage-preserving TRN of TFs, there may exist specific gatekeeper TFs which play such a critical role that they act as a molecular lynchpin, and their sole inhibition may singlehandedly permit cell reprogramming.

For a factor to be a true “Gatekeeper” Transcription Factor (GTF), its singular targeted inhibition should be capable of reversing lineage preservation, commitment and specification and restoring multipotent and/or pluripotent differentiation capability. Furthermore, the targeted siRNA-mediated inhibition of the identified GTF in differentiated cells should be a viable alternative to the forced overexpression of pluripotency factors in such cells as a means to generate multipotent and/or pluripotent stem cells from them without the need for permanent genetic modification. We hypothesized that true GTFs do exist whose physiologic expression is critically necessary to inhibit cell plasticity and pluripotency and safeguard the processes of lineage specification, commitment and preservation in differentiated cells.

## RESULTS

To test our hypothesis, we decided to examine the process of mesoderm specification, commitment and preservation that occurs in ESCs to give rise to the prototypical differentiated mesenchymal cells like fibroblasts as a model system. To identify the most critical lineage-defining TFs of Rat Embryonic Fibroblasts (REFs), we pursued an unbiased and comprehensive experimental discovery approach involving generation of stable intra-species somatic cell synkaryon hybrids (HFs) (an accepted model of gene silencing of TFs responsible for preservation of parental cell identity that are mononuclear, tetraploid with practically no chromosome loss and stable genotype) created by fusing REFs of mesenchymal origin with rat hepatoma (RH) cells of endodermal origin after genetically engineering each cell line so that each possessed a unique yet separate xenobiotic resistance mechanism (Bulla et al., 2010).

Parental mesenchymal and endodermal cells were incapable of surviving exposure to either one or the other xenobiotic, and perished. The cells that survived exposure to both xenobiotics were true synkaryon hybrids wherein the obligate gene silencing of mesenchymal and endodermal master regulatory lineage-instructive TFs displayed by such cells permits other *trans* regulatory elements of both to coexist in complementary fashion. By subjecting the starting parental as well as resultant hybrid cells to genome-wide transcriptomic profiling, we identified pivotal master regulatory TFs which are not only the most critical for mesenchymal lineage commitment and preservation but whose downregulation constitutes an imperative requirement for reacquisition of cellular plasticity by these cells. First, we selected 149 TFs that demonstrated greater than 5-fold overexpression in REFs compared to RH cells and referred to these as mesenchymal fibroblast-enriched TFs (MFEFs) [Fig. 1A]. We then screened these MFEFs to identify 86 TFs that demonstrated greater than 2.5-fold repression in hybrids compared to REFs and referred to these as mesenchymal fibroblast-specific TFs (MFSFs) [Fig. 1A]. Snai2 and Prrx1 were the top MFSFs that were repressed in hybrids compared to REFs [Fig. 1D], and were the candidate gatekeeper TF master regulators of mesenchymal fate that were selected for further investigation.

**Fig. 1.**
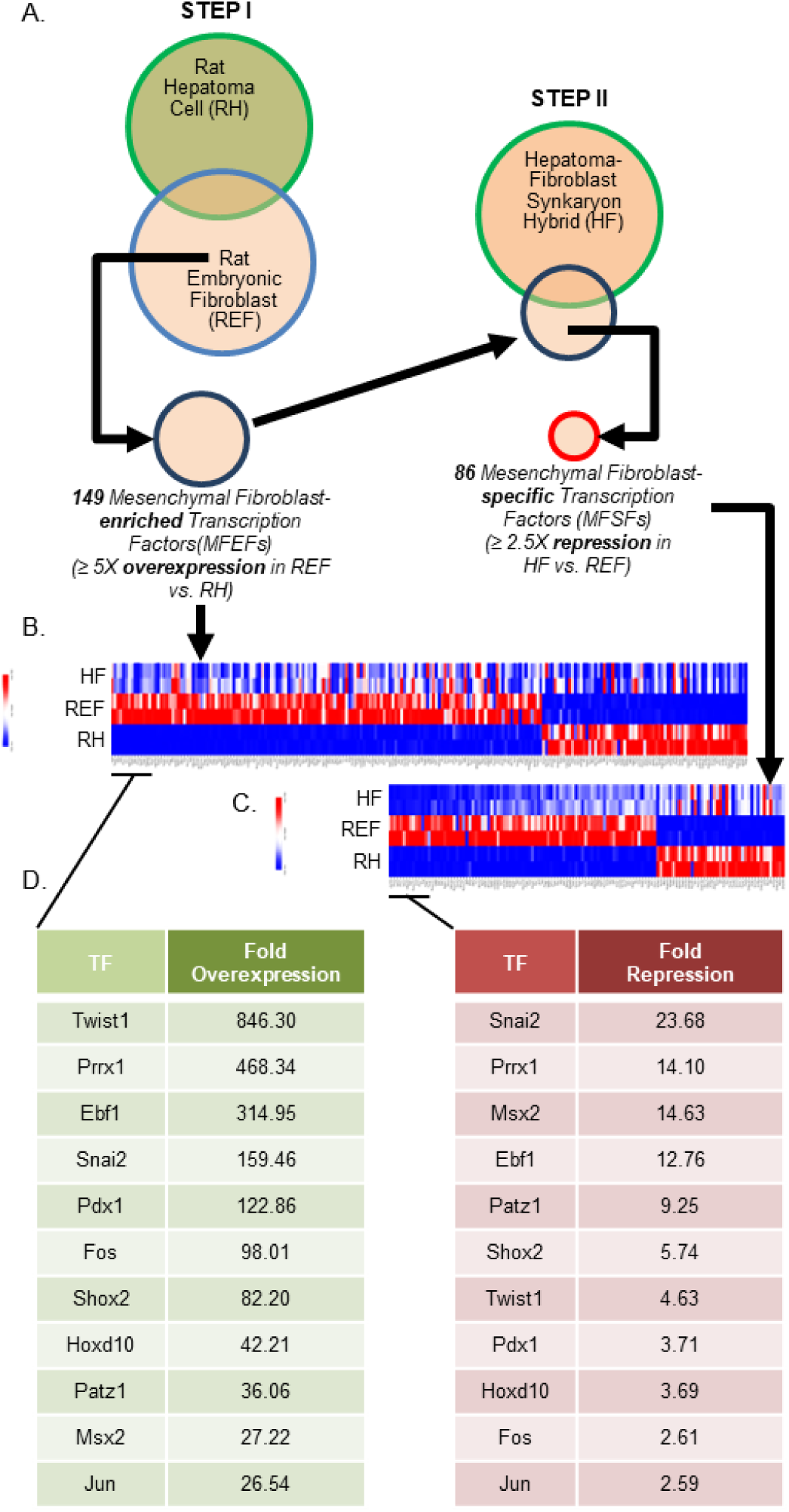
Identification of core transcriptional regulators that define mesenchymal fibroblast identity. A) Venn diagram representation of the two-step bioinformatic strategy to identify core transcriptional regulators of mesenchymal fibroblast cell identity. By selecting transcription factors (TFs) that displayed a > 5-fold expression in rat embryonic fibroblast (REF) cells of mesenchymal origin when compared to rat hepatoma cell (RH) of endodermal origin, 149 TFs were grouped as mesenchymal fibroblast-enriched TFs (MFEFs). Out of these 149 MFEFs, 86 TFs that displayed >2.5-fold repression in the hepato-fibroblast synkaryon cell hybrid (HF) when compared to REFs, were identified as mesenchymal fibroblast-specific TFs (MFSFs). B) Gene expression microarray heatmap of REF, RH, and HF cell lines showing all genes that displayed >5fold overexpression in REF compared to RH cells. TF gene-members were shortlisted as MFEFs. C) Gene expression microarray heatmap of HF and REF cell lines showing all genes that displayed >2.5fold repression in HF when compared to REFs. TF gene-members were identified as MFSFs. D) List of top ten MFEFs and MFSFs are tabulated. It is interesting to note how the rank order of the top MFEFs changed in the core MFSFs list, after applying the two-step selection strategy.

We next validated expression of the candidate MFSFs identified from our microarray analysis by quantitative reverse transcriptase polymerase chain reaction (q-RTPCR) and confirmed repression of fibroblast specific genes (Prrx1, Snai2 and Col1a1) in somatic cell hybrids [Fig. 2C]. To ensure the robustness of these identified MFSFs we ectopically overexpressed the top two MFSFs (Snai2 and Prrx1) individually in the somatic cell hybrids. Generation of G418 resistant Snai2 and Prrx1 overexpressed hybrid clones, respectively, led to re-acquisition of fibroblast traits (morphology, structure and functionality) in the hybrid cells. Snai2 and Prrx1 clones attained fibroblast-like spindle-shaped morphology [Fig. 2A,B], re-expressed fibroblast prototypical gene Col1a1 as confirmed by IF and q-RTPCR [Fig. 2B,C] and displayed significantly increased migration ability compared to the HF hybrids especially in response to TGF-Beta stimulation, a characteristic fibroblast-specific functional trait (Acharya et al., 2008) [Fig. 2D], while individual siRNA-mediated knockdown of Snai2 and Prrx1 in REFs resulted in loss of spindle-shaped morphology, loss of Col1a1 expression and diminished migratory capacity compared to parental REFs.

**Fig. 2.**
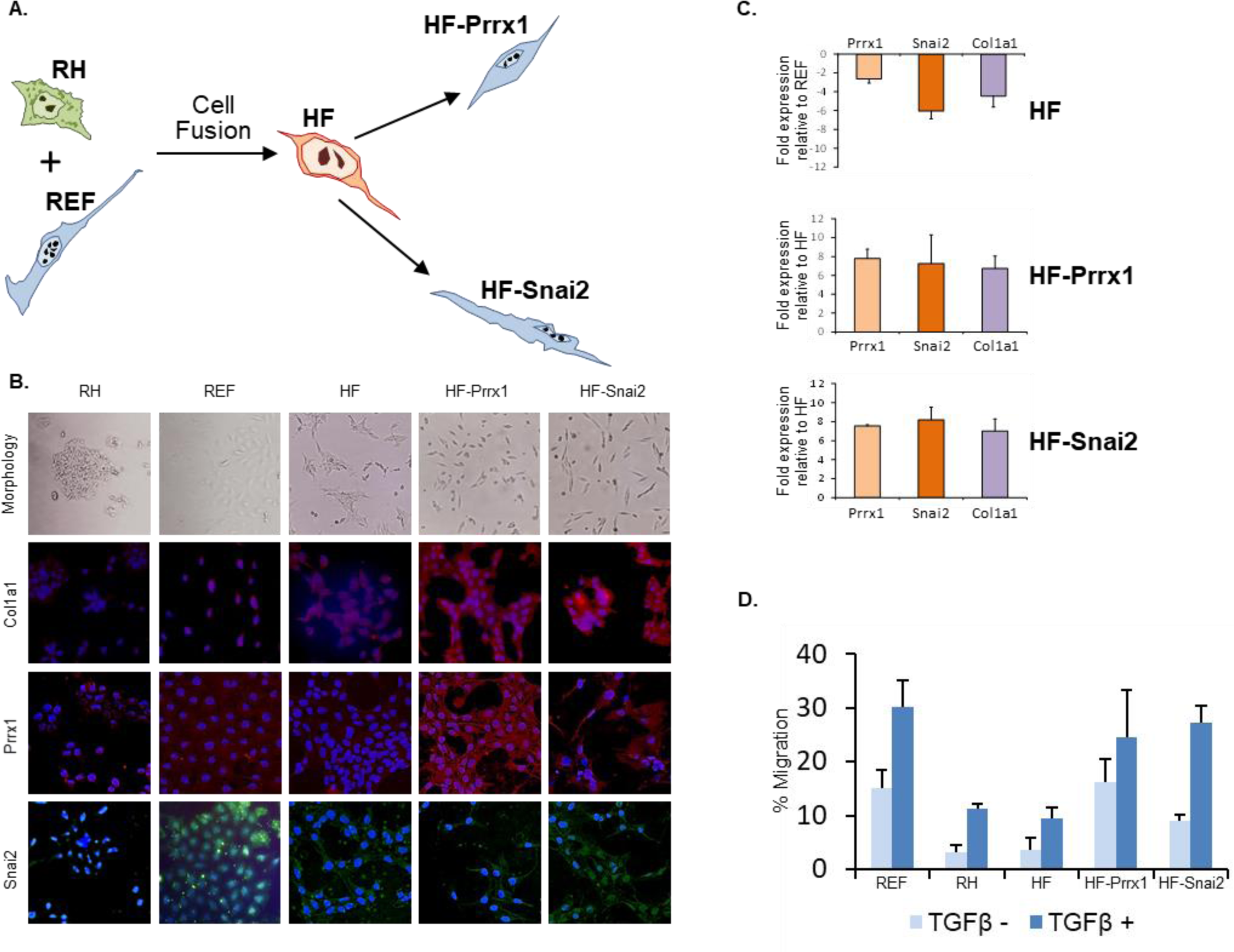
Effect of engineered re-expression of Prrx1 and Snai2 in HF cells. A) Schematic diagram depicting generation of stable intra-species hepatoma fibroblast (HF) hybrid by fusion of rat hepatoma (RH) and rat embryonic fibroblasts (REFs). Generation of HF-Prrx1 and HF-Snai2 clones by ectopic expression of Prrx1 &Snai2 in HF. B) Row 1: Photomicrographs shows round, spindle shaped and irregular morphology of RH, REF and HF cells respectively, re-acquisition of fibroblast-like morphology in HF-Prrx1 and HF-Snai2 clones. Row 2,3&4: IF pictures showing expression of prototypical mesenchyme-fibroblast markers Col1a1, Prrx1 & Snai2 in RH, REF, HF, HF-Prrx1and HF-Snai2 clones respectively. (Col1a1and Prrx1 are stained in red, Snai2 is stained in green and DAPI in blue) (Scale bar, 10µm). C) Fold repression of Prrx1, Snai2 and Col1a1 in HF hybrids vs REFs validated by qRTPCR (Top), re-establishment of Prrx1, Snai2 & Col1a1 expression in HF-Prrx1 and HF-Snai2 clones (middle & bottom, respectively). D) Bar graph depiction of TGF Beta-responsive cell migration ability of RH, REF, HF, HF-Prrx1 and HF-Snai2 clones.

After confirming the pivotal importance of Snai2 and Prrx1 in preserving the structural and functional cellular identity of mesenchyme derived REFs, we next set out to determine whether the individual forced transient repression of either of these two critical TFs (utilizing a transfection protocol with siRNAs specific for either Snai2 or Prrx1) could lift the cell fate restrictions tied to mesenchymal and fibroblast identity permitting acquisition of cellular plasticity, and therefore render them prone to cellular reprogramming. In order to test our hypothesis, we incubated the siSnai2REF and siPrrx1REF cells in induction media of alternate lineage cells of mesenchymal origin. When cultured in adipogenic media for 14 days, siSnai2REF and siPrrx1REF cells both attained lipid droplet-filled spherical morphology akin to adipocyte cells. Lipid vacuoles were visualized by light microscopy, stained red with Oil Red O [Fig. 3A,D] and these cells expressed adipocyte lineage-appropriate TF Cebpa as evaluated by q-RTPCR [Fig. 3G] and immunofluorescence (IF) [Fig. 3A]. Similarly, individual knockdown of Snai2 or Prrx1 in the parental REF by siRNA transfection and maintenance of the siSnai2REF and siPrrx1REF knockdown cells in either osteogenic or chondrogenic media for 14 days resulted in functional osteocytes and chondrocytes, respectively, for both types of knockdown cells. Lineage-appropriate function of forming bone was evidenced by positive staining with Alizarin Red [Fig. 3B,E] while cartilage formation was confirmed by positive staining with Alcian blue, [Fig. 3C,F]. Expression of osteocyte and chondrocyte lineage-instructive TFs Runx2 and Sox9, respectively, was also confirmed using q-RTPCR [Fig. 3H,I] and immunofluorescence (IF) [Fig. 3B,C].

**Fig. 3.**
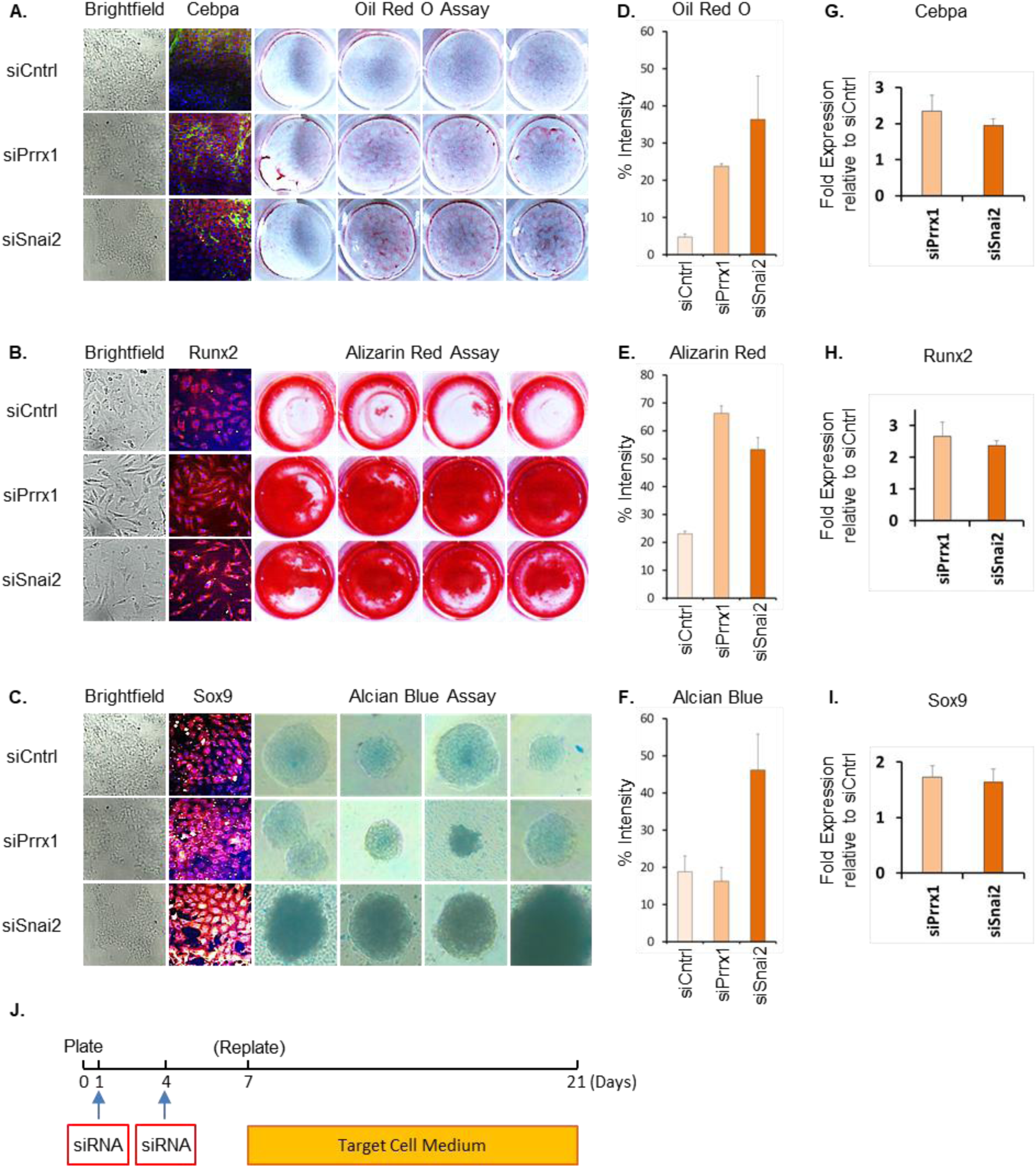
siRNA mediated transient repression of Snai2 or Prrx1 in REFs induces ex vivo adipocytic, osteocytic or chondrocytic transdifferentiation. A, B & C) Brightfield photomicrographs of REFs transfected with siCntrl, siPrrx1 & siSnai2, (Rows 1,2 and 3) that were then incubated in adipogenic media, osteogenic media and chondrogenic media respectively for 14 days. IF photomicrographs assess expression of adipogenic TF Cebpa (Red), osteogenic TF Runx2 (Red) and chondrogenic TF Sox9 (Red) in all 3 groups with DAPI nuclear counterstain in Blue. (Scale bar, 10µm). D,E &F) % intensity of Oil Red O, Alizarin Red and Alcian Blue staining to ascertain adipogenesis, osteogenesis and chondrogenesis as quantitated using Image J software analysis. G,H&I) q-RTPCR analysis to assess increase in relative expression of Cebpa, Runx2 and Sox9 compared to housekeeping genes (Ppia, Spp1 and Ppia) in siPrrx1 and siSnai2 treated REFs versus siCntrl treated REF cells when incubated in adipogenic, osteogenic and chondrogenic media for 14 days respectively. Data are represented as mean±SD of 3 independent experiments. J) Schematic diagram illustrating the process of siRNA-mediated cellular reprogramming in the absence of exogenous factors.

Next, we sought to determine whether the siRNA-mediated transient repression of either Snai2 or Prrx1 and incubation in suitable cell culture conditions could effectively transform parental fibroblasts into a more primitive precursor cell state, developmentally precedent to the mesenchymal differentiated state. siSnai2REF and siPrrx1REF cells when cultured in rat mesenchymal stem cell (rMSC) media for 14 days resulted in transformation to dedifferentiated multipotent stem-like cells (dMSCs), with altered morphology and enhanced expression of the prototypical MSC TF Myc (validated by both IF [Fig. 4A] and qRTPCR [Fig. 4C]). dMSCs showed significantly enhanced expression of alkaline phosphatase, an indicator of undifferentiated stem cell activity [Fig. 4B]. When siSnai2REF and siPrrx1REF cells were cultured over mitomycin-C inactivated REFs in rat embryonic stem cell (rESC) media (with 2i/LIF) for 14 days (Jackson et al., 2010), attainment of dedifferentiated pluripotent stem cell traits occurred only in the siSnai2REF group [Fig. 4D] with expression of characteristic pluripotency factors like Sox2, Klf4 and Nanog, [Fig. 4E,F]. Since these cells were obtained by lifting reprogramming barriers (lineage preserving TFs) that impede dedifferentiation, we refer to these cells as dedifferentiated pluripotent stem-like cells (dPSCs). Suspension culture of dPSCs, when placed in rESC differentiation media for 8 days, formed embryoid bodies that comprised of all three germ layers and stained positive for triple germ layer-specific TFs namely Sox2 and Otx2 (ectoderm) [Fig. 4F], Brachyury and Hand1 (mesoderm) [Fig.4G] and Gata4 and Sox17(endoderm) [Fig. 4H] as validated by IF staining and confocal microscopy.

**Fig. 4.**
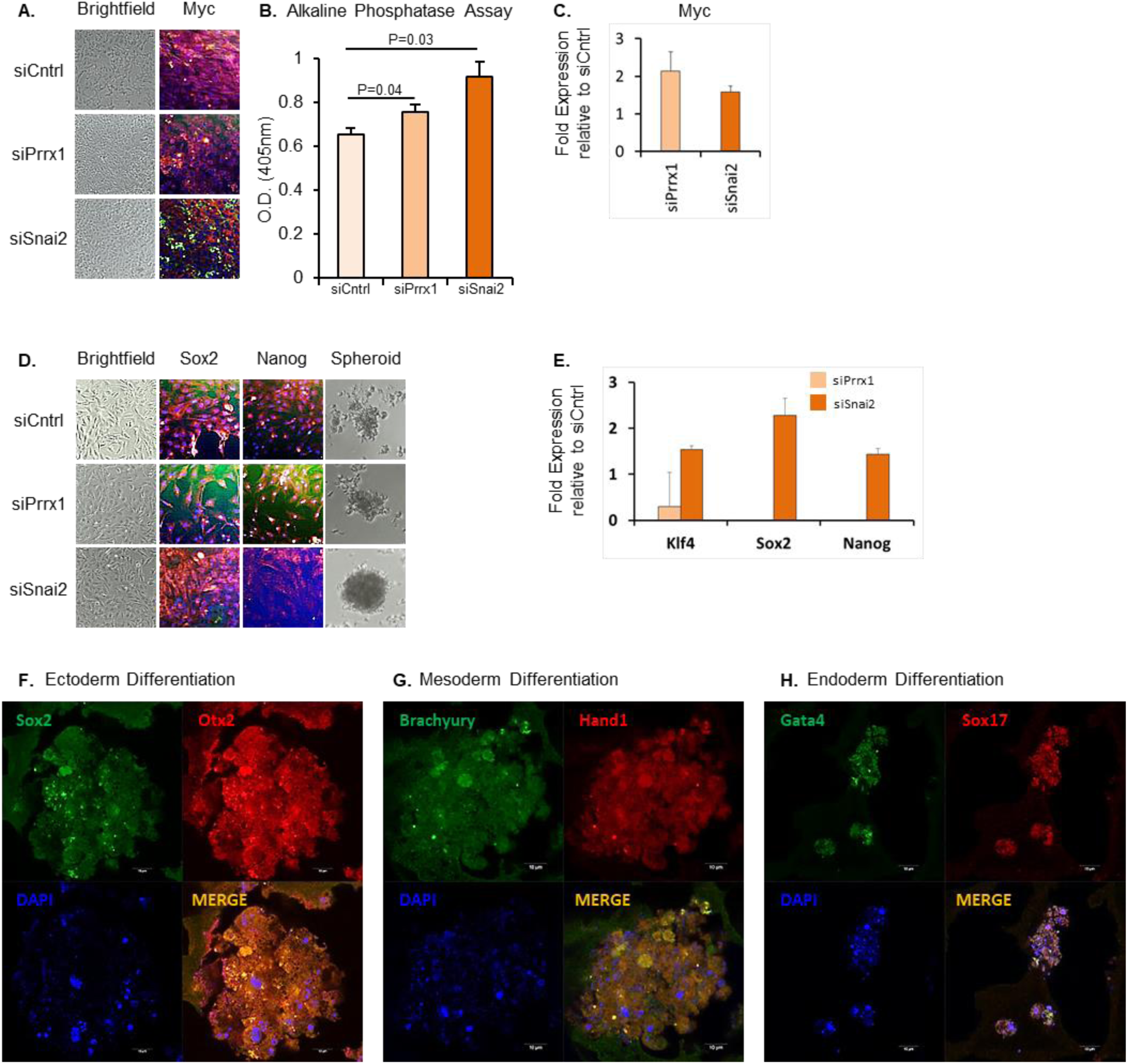
Dedifferentiation of Rat Embryonic Fibroblasts to dMSCs and dPSCs by transient repression of Snai2 and Prrx1. A,D) Brightfield photomicrographs of REFs transfected with siCntrl, siPrrx1 & siSnai2, (Rows 1,2 and 3) that were incubated in MSC media or Rat ESC media for 14 days. IF photomicrographs assess expression of MSC TF Myc (Red) and ESC TFs (Sox2 and Nanog in Red) and DAPI (Blue) in all three groups (Scale bar, 10µm). D) (Column 4) Brightfield photomicrographs of free-floating embryoid bodies on day 8 of suspension culture in differentiation media following transfection of REFs with siCntrl, siPrrx1 & siSnai2 and incubation in Rat ESC media for 14 days. B) MSC activity was ascertained using Alkaline Phosphatase Assay. C,E) q-RTPCR analysis to assess increase in relative expression of Myc, Klf4, Sox2 and Nanog compared to housekeeping gene (B2m) in siPrrx1 and siSnai2 treated REFs when incubated in MSC and ESC media respectively for 14 days. F, G and H) IF Confocal microscopy images assess expression of Ectodermal TFs Sox2 (green) and Otx-2 (red), Mesodermal TFs Brachyury (green) and Hand1 (red), and Endodermal TFs Gata-4 (green) and Sox17 (red) of embryoid bodies at Day 8 in the siSnai2 group, nuclear counterstain DAPI (blue).

## Discussion

Lineage specification and cell fate decision is enforced by the expression of cell type-specific TFs and the underlying epigenetic state. These factors act as powerful safeguards against acquisition of cellular plasticity and subsequent alteration of the cell’s lineage commitment and identity. In order to understand the role and extent of regulation that these lineage instructive TFs exert on deciding and preserving lineage commitment and cell fate, we first identified the single most pivotal GTFs from among the multiple TFs that comprise the TRN of REFs, our model system. We then transiently repressed the identified GTFs in REFs to determine whether this alone, in the absence of exogenous TFs or small molecule cocktails, was sufficient to affect a switch between a state of lineage commitment to a state of cell plasticity. The targeted inhibition of single GTFs proved successful in disrupting the entire lineage preserving machinery of a differentiated cell type (REFs) thereby rendering the cells plastic and ready for a reprogramming cascade.

Transient inhibition of cell lineage-specific TFs is thus an independent mechanism by which a cell can be reprogrammed in the absence of exogenous TFs or small molecule cocktails. Though TF-mediated somatic cell reprogramming (whereby target cell-specific TFs are ectopically overexpressed) and small molecule driven reprogramming (whereby pathway modulators induce changes in the epigenetic landscapes) are powerful cell reprogramming tools employed to generate pluripotent cells or differentiated cells of alternate desired lineages from lineage-committed cells, the efficiency of such processes is impaired by the persistence of the starting cell’s TRN and residual epigenetic memory (Nashun et al., 2015). Studies have shown that in intermediately transformed cells, some specific genes of the starting cells are persistently expressed, whose siRNA mediated knockdown considerably improved reprogramming efficiency (Ebrahimi, 2015; Mikkelsen et al., 2008). Though multiple gene knockdown of the starting fibroblast cell’s Transcriptional Regulatory Network (TRN) yielded differentiated adipocytes (Tomaru et al., 2014) this concept however was never quite extended to deciphering the impact of repression of a single critical lineage-instructive TF of the starting cell in affecting transdifferentiation. Similarly, while TFs that interfere with dedifferentiation have been demonstrated to induce cell-type-specific transcriptional profiles, repression of the identified TFs in prior studies were not successful in permitting dedifferentiation (Hikichi et al., 2013), likely because of differences in the employed siRNA transfection protocol. It was noted that Snai2 was among the earliest genes to be repressed during traditional OSKM mediated somatic reprogramming efforts (Cacchiarelli et al., 2015; Mikkelsen et al., 2008; Polo et al., 2012) and that induced repression of Prrx1 or Snai2 enhanced both OSKM (Yang et al., 2011) and Nanog (Gingold et al., 2014) mediated fibroblast reprogramming, respectively. However, to our knowledge, there are no prior studies that successfully identify pivotal cell-specific master regulators of cell fate whose upregulation is critically necessary for lineage commitment, specification and identity preservation and whose downregulation, by the same token, is imperatively necessary for acquisition of cellular plasticity adequate to permit either transdifferentiation to alternate cell fates, or dedifferentiation to multipotent/pluripotent primitive cell states, in the absence of any exogenous transdifferentiation or pluripotency TFs or pathway modulatory cocktails.

Using REFs as a model system, our experimental results show that transient repression of a single pivotal GTF of REFs (Snai2 or Prrx1 in our case) is adequate to overcome the lineage preservation effect of the starting cells’ TRN, and is complete enough to permit acquisition of cellular plasticity and overcome barriers to nuclear reprogramming. Furthermore, when incubated in appropriate media, the transient repression of the identified pivotal master regulators Snai2 and Prrx1 in fibroblasts can lead to direct lineage conversion of the starting cell to alternate cell types (such as adipocytes, chondrocytes or osteocytes). While repression of either Prrx1 or Snai2 was capable of dedifferentiating REFs to multipotent dMSCs, siSnai2 demonstrated the ability to dedifferentiate REFs to dPSCs. We surmise that this is perhaps due to Snai2 first exerting transcriptional regulatory control in Day 3.5 CD326^-^CD56^+^ embryonic mesodermal precursor cells (160X higher expression than ESCs), which precedes the first expression of Prrx1, in the hierarchy of mesenchymal cell development (Evseenko et al., 2010). Our described method resets a cell to an earlier point in its developmental timeline, likely derepressing plasticity maintenance factors, enabling its innate differentiation mechanism to then proceed as guided by components provided in its growth medium.

The detailed molecular mechanisms underlying the cellular transformations brought about by transient repression of Snai2 and Prrx1 remain unknown as do any potential chromatin landscape alterations that may have resulted from such RNAi manipulation. In conclusion, these results expand our molecular understanding of lineage commitment, specification and fate preservation and reveal that certain pivotal gatekeeper TFs that are master regulators of cell fate can also be considered to constitute a molecular Achilles’ heel in a specific cell’s identity. They act as the molecular lynchpin of cell fate preservation and may possibly be the single most important reason why cell identity is not immutable. If gatekeeper TFs of a specific cell type can be correctly identified, the siRNA mediated cell-specific transient repression of such gatekeeper TFs can be a tenable strategy to help achieve cell identity switch of such cells to alternate transdifferentiated cell states or to dedifferentiated multipotent/pluripotent cell states, without the need for permanent genetic modification of the targeted cells. Additionally, this may potentially enhance reprogramming efficiencies of exogenous TF/small molecule driven protocols both in terms of cell yield and process time and eventually help realize the goal of reliably obtaining clinically relevant somatic cells or multipotent/pluripotent stem cells for therapeutic use.

## Acknowledgements

Funding for this study was provided by Eastern Illinois University Graduate Academic Award (TR), National Science Foundation Grant 0841653 (GAB), Recode Biosciences Research Grant SC010116 (PSR).

## Author Contributions

Conceptualization: TR, PSR; Data Curation: TR, AS, PSR; Formal Analysis: TR, PSR; Funding Acquisition: TR, GAB, PSR; Investigation: TR, AS; Methodology: TR, GAB, PSR; Project Administration: GAB, PSR; Resources: GAB, PSR, Supervision: GAB, PSR; Validation: GAB, PSR; Visualization: TR, PSR; Writing – original draft: TR, PSR; Writing – review and editing: TR, AS, GAB, PSR.

## Methods

### Cell lines and Culture Conditions

The rat hepatoma cell line FT02B is an ouabain-resistant, thymidine kinase TK-deficient H4IIEC3 derived clone(23). RAT-1 is a SV40-transformed rat embryonic fibroblast (REF) cell line that expresses functional thymidine kinase (Botchan et al., 1976). HF is a hybrid cell line generated by the fusion of FT02B and RAT-1 cells (Bulla et al., 2010). All cells were maintained in 1:1 Ham’s F12 Dulbecco’s modified Eagle’s medium (FDV) containing 5% fetal bovine serum (FBS) (Thermo Fisher Scientific) and 5ml/500ml 100X penicillin-streptomycin (Thermo Fisher Scientific) at 37°C in a 5% CO2 incubator.

### Microarray Analysis

RNA was extracted using RNA Easy Mini Kit (Qiagen) according to manufacturer’s protocol. RNA purity and concentration were determined using an Epoch Nano-drop spectrophotometer at 260 and 280 nm. Following extraction, the RNA samples from all cell lines were sent to the W.M. Keck Center for Comparative and Functional Genomics in the Roy J. Carver Biotechnology Center at the University of Illinois in Urbana-Champaign for Microarray Analysis. RNA was hybridized to Affymetrix GeneChip® Rat Genome 230 2.0 Expression Arrays (Affymetrix, Inc., Santa Clara, CA) utilizing the GeneChip (Expression 3’ Amplification One-Cycle Target Labeling and Control Reagents kit according to the manufacturer’s instructions). The chips were scanned with a GeneChip Scanner model 3000 7G Plus. Gene expression was detected and normalized.

### cDNA synthesis and q-RTPCR Analysis

RNA was extracted using RNA Easy Mini Kit (Qiagen) according to manufacturer’s protocol. RNA purity and concentration were determined using an Epoch Nano-drop spectrophotometer at 260 and 280 nm. Reverse transcription was performed with the Superscript III First Strand cDNA Synthesis kit (Invitrogen) following manufacturers protocol using a Thermal Cycler (Applied Biosystems™, USA). The resulting cDNA was used for Quantitative real-time RTPCR (qRTPCR) performed in 20μl reaction mixture [2µl of cDNA (5ng/µl), 6.75µl of sterile nuclease free water, 10µl of Fast SYBR® Green Master Mix (Applied Biosystems™, USA) and 1.25µl of gene specific primer (0.5µm from IDTDNA)] on a Step One Plus Real-Time PCR System (Applied Biosystems™, USA). PCR amplification was carried out at 95°C for 10s, followed by 40 cycles at 95°C for 5s and at 60°C for 30s. The amplification signal of the target genes’ mRNA was normalized against endogenous house-keeping genes’ mRNA in the same reaction. The list of primers used for q-RT-PCR have been provided in the Key Resources Table.

### Overexpression of Candidate Genes (Prrx1 and Snai2) in the HF Hybrids

Expression vectors containing full length rat genes Snai2 and Prrx1 (Origene, Inc.) were introduced into the hybrid cell line HF by lipofection using Lipofectamine Plus reagent (Invitrogen, Inc.) according to manufacturer’s protocol. After 2–3 weeks, stable G418 resistant clones were selected, expanded, and used for subsequent experiments.

### Knockdown of Candidate Genes (Prrx1 and Snai2) in the REF Cells

RAT-2 REFs (acquired from ATCC), were seeded (four replicates) in 6-well cell culture plates (Nunc) at a density of 2 × 106 cells/well and incubated at 37°C in a CO2 incubator one day before transfection was performed with 50 nM (final concentration) of pooled small interfering RNA (siRNA, Santa Cruz Biotech), 2.5μl of Lipofectamine RNAiMAX (Invitrogen, USA) and Opti-MEM (Invitrogen, USA) according to the manufacturer’s instructions. RNAi negative universal control MED (Invitrogen) was used to calibrate siRNA transfection. Repeat transfection was performed on Day 4, in order to achieve sustained inhibition, prior to using cells for subsequent experiments on Day 7 (Fig3J).

### Cell Migration Assay

Trans-well cell migration assay for REFs, HFs, HF-Snai2 and HF-Prrx1 cells was carried out using 24-well cell migration kit from Trevigen (Catalog# 3465-024-K) following manufacturers protocol. The assay plate was read using a Biotek Synergy HT plate reader at 485 nm excitation, 520 nm emission. The experimental data, in terms of relative fluorescence units (RFU) was converted into cell numbers to determine the number of cells that have migrated.

### Adipogenic induction

REFs were cultured in a DMEM media with 10% FBS and 1% Pen-Strep. When the cells reached 70% confluency, they were trypsinized, pelleted and counted. 30000 cells were plated in 4 wells for each experimental group (siCntrlREF, siPrrx1REF and siSnai2REF) in a 96 well plate. The wells were transfected with control, Prrx1 and Snai2 siRNA. Seven days post initial siRNA transfection, transfected cells were treated with adipogenic induction medium (A) for three days (Cyagen Biosciences, Cat# GUXMX-90031) followed by maintenance medium (B) for 11days. On the 14th day, oil red staining was performed to assess adipogenic activity and cell pellets were preserved for further downstream analyses.

### Oil Red O staining

Two weeks post adipogenic induction, cells were rinsed with phosphate buffered saline (PBS) and fixed with 4% paraformaldehyde (Sigma Aldrich) in PBS. The formaldehyde solution was removed by tilting the plate and rinsed with sterile water. Each well was then covered with 60% of isopropanol and incubated for 5 minutes. Isopropanol solution was pipetted out and 2 ml of working solution of Oil Red O (Sigma Cat# O1391) was added in each well and incubated for 20 minutes at room temperature. Wells were then washed with water until the water ran clear. All wells were kept wet with water and viewed under the microscope. Lipid droplets were stained red with Oil Red O. To record the OD readings Isopropanol solution was pipetted into a fresh 96 wells microtiter plate (Corning NBS Microplate) and OD reading was recorded at 490/500/510 nm, respectively Biotek Synergy HT plate reader.

### Osteogenic induction

REFs were cultured in a DMEM media with 10% FBS and 1% Pen-Strep. When the cells reached 70% confluency, they were trypsinized, pelleted and counted. 30000 cells were plated in 4 wells for each experimental group (siCntrlREF, siPrrx1REF and siSnai2REF) in a 96 well plate. The wells were transfected with control, Prrx1 and Snai2 siRNA. Seven days post initial siRNA transfection, transfected cells were placed in osteogenic medium (DMEM, FBS 10%, 50 µg/ml ascorbic acid, 10mM β-glycerophosphate, 10nM dexamethasone). Every 3 days old media was withdrawn and new Osteogenic media was added. After 14 days, cells were washed in cold PBS, fixed with 4% PFA in PBS and stained with 40mM Alizarin Red that stained for Calcium to ascertain osteogenic activity and cell pellets were preserved for further downstream analyses.

### Alizarin Red staining

Two weeks post osteogenic induction, cells were rinsed with phosphate buffered saline (PBS) thrice and fixed with 4% paraformaldehyde (Sigma Aldrich) in PBS for 15min at room temperature. The formaldehyde solution was removed, and the cells rinsed with sterile diH2O. Water was removed completely and 1mL of 40mM Alizarin Red S (ARS) was added and alizarin red staining was carried out as per manufacturer’s protocol (ScienCell, ARS Staining Quantification Assay ARed-Q Catalog #8678).

### Chondrogenic induction

REFs were cultured in a DMEM media with 10% FBS and 1% Pen-Strep. When the cells reached 70% confluency, they were trypsinized, pelleted and counted. 30000 cells were plated in 4 wells for each experimental group (siCntrlREF, siPrrx1REF and siSnai2REF) in a 96 well plate. The wells were transfected with control, Prrx1 and Snai2 siRNA. Seven days post initial siRNA transfection, transfected cells were placed in chondrogenic medium (MSC go Chondrogenic XF™, Biological Industries) and incubated for 14 days in incubator (37°C, 5% CO2). The chondrogenic medium was changed every 3 days for 14 days following which Alcian Blue staining was used to evaluate chondrogenic activity and cell pellets were preserved for further downstream analyses.

### Alcian Blue staining

Two weeks post osteogenic induction, cells were rinsed with phosphate buffered saline (PBS) thrice and fixed with 4% paraformaldehyde (Sigma Aldrich) in PBS for 15min at room temperature. The formaldehyde solution was removed, and the cells rinsed with sterile diH2O. Water was removed completely and 0.2ml of 1% Alcian Blue solution (Sigma; A-3157) was added to each well and alcian blue staining carried out as per kit manufacturers protocol (MSC go Chondrogenic XF™, Biological Industries).

### Generation of dMSCs

REFs were cultured in a DMEM media with 10% FBS and 1% Pen-Strep. When the cells reached 70% confluency, they were trypsinized, pelleted and counted. 30000 cells were plated in 4 wells for each experimental group (siCntrlREF, siPrrx1REF and siSnai2REF) in a 96 well plate. The wells were transfected with control, Prrx1 and Snai2 siRNA. Seven days post initial siRNA transfection, transfected cells were placed in Rat MSC medium (Rat MSC growth medium kit, Cell Applications, Inc.) and incubated for 14 days in incubator (37°C, 5% CO2). The MSC medium was changed every 3 days for 14 days following which Alkaline Phosphatase Assay (Anaspec EGT, Catalog # AS-72146) was used to assess mesenchymal stem cell activity according to manufacturer’s instructions and cell pellets were preserved for further downstream analyses.

### Generation of dPSCs and Embryoid bodies

All three groups (siCntrlREF, siPrrx1REF siSnai2REF) of rat embryonic fibroblasts (REF) cells were transferred (seven days post initial siRNA tranfection) onto REF feeder layer inactivated with mitomycin C (Sigma) in replicates of four in a 12 well plate and maintained in ESC media (DMEM/F12 supplemented with 20% knockout serum replacement (KSR) and 8ng/ mL of fibroblast growth factor (FGF-2) for 14days in incubator (37°C, 5% CO2)(Gomes et al., 2010). After 14 days, the generated dPSCs were transferred to non-adherent dishes and cultured in suspension culture for 8 days in differentiation media to induce Embryoid Body (EB) formation. Cell pellets from matching experiments were preserved for further downstream analyses. dPSCs were differentiated into each of the three germ layers according to the Human Pluripotent Stem Cell Functional Identification Kit (R&D Systems®, Catalog # SC027B).

### Fixation and Cryopreservation of Embryoid Bodies

The EBs were collected from the culture dish with a pipette and transferred to a 15 mL conical tube until all material sank to the bottom of the tube. After gently removing the media, the EBs were fixed with a 4% paraformaldehyde (PFA) solution in PBS for 30 minutes in room temperature. Next the PFA solution was removed and the cells washed with PBS for 5 min.

### Immunofluorescence Staining

Immunofluorescence staining for FT02B, REF, HFs, HF-Prrx1and HF-Snai2 cells was performed following standard protocol using appropriate monoclonal primary antibody [Prrx1(Novus Biologicals), Snai2 (SantaCruz Biotechnologies), Col1a1 (Origene), Myc (SantaCruz Biotechnologies), Sox2 (Invitrogen USA), Nanog (Invitrogen USA)] used at concentrations of 5ug/ml], followed by labeling with Alexafluor-conjugated secondary antibody (Abcam) (1:100 dilution). Samples were subsequently mounted in Prolong Gold antifade mounting media (Thermo Fisher Scientific) and analyzed with a fluorescent microscope (Leica DMIL LED) using 20X magnification and appropriate wavelength settings.

### Immunofluorescence detection of three germ layers

The fixed samples were embedded in agarose to trap and immobilize the embryoids. Once the agarose solidified, samples were kept at 4°C until further processing. The samples were embedded in paraffin wax and sectioned at 5-6µ thickness using a microtome. The sections were then adhered to positively charged slides and then dewaxed in Histoclear and rehydrated as per protocol(Sivaguru et al., 2013). The rehydrated slides were permeabilized with 0.5% TritonX-100 in phosphate buffered saline and labeled with human three germ layer 3-color immunocytochemistry kit from R&D Biosystems following the manufacturer protocol (Catalog# SC022, R and D systems, Inc., MN, USA). Briefly, the sections were blocked by 10% normal donkey serum (Jackson Laboratories, USA) containing 1% BSA (Thermofisher, USA). After blocking the sections were incubated with a combination of two conjugated primary antibodies for each lineage of interest such as, ectoderm (Anti-Sox1 conjugated with Northern Lights 493 and anti-Otx-2 conjugated with Northern Lights 557), mesoderm (Anti-Brachyury conjugated with Northern Lights 557 and anti-Hand1 conjugated with Northern Lights 637), and endoderm (Anti-Gata-4 conjugated with Northern Lights 493 and anti-Sox17 conjugated with Northern Lights 637). Each combination was incubated for 3h at room temperature in the dark, washed with PBS containing BSA, counterstained with Hoechst nuclear dye (Thermofisher, USA), mounted in Prolong Gold antifade mounting media (Thermofisher, USA), left for 24h at room temp and stored in 4°C after sealing with nail polish.

### Airyscan Super-Resolution Imaging

The labeled sections were imaged under a Zeiss LSM 880 Laser Scanning Microscope with Airyscan Super-Resolution system. Excitation and emission wavelengths that were collected include: 405 nm excitation (emission collected between 410-460 nm), 488 nm excitation (emission collected between 500-550nm), 561 nm excitation (emission collected between 570-615 nm) and 633 nm excitation (emission collected between 650-700 nm). An internal GaAsP photo-multiplier tube detector was used to scan entire stone thin sections using a 63x Plan Apochromat (NA 1.4 oil immersion objective. Several images were taken at different sections and locations for each sample. The images were recorded at a super-resolution sampling frequency of 40nm per pixel. The raw data were processed for super-resolution in the same program Zeiss Zen software and pseudo-colored for visualizing four channels.

### Statistics and Reproducibility

All in vitro transdifferentiation and reprogramming related experiments were repeated independently at least three times with similar results. Replicates from failed experiments due to technical issues were excluded from analysis. Details on sample sizes and reproducibility are in the figure legends. Wherever appropriate, data are presented as the mean ± s.e.m. Wherever appropriate, comparisons between groups used the Student’s t-test assuming two-tailed distributions.

### Data Availability

All data supporting the findings of this study are available from the corresponding author on reasonable request.

## Key Resources Table

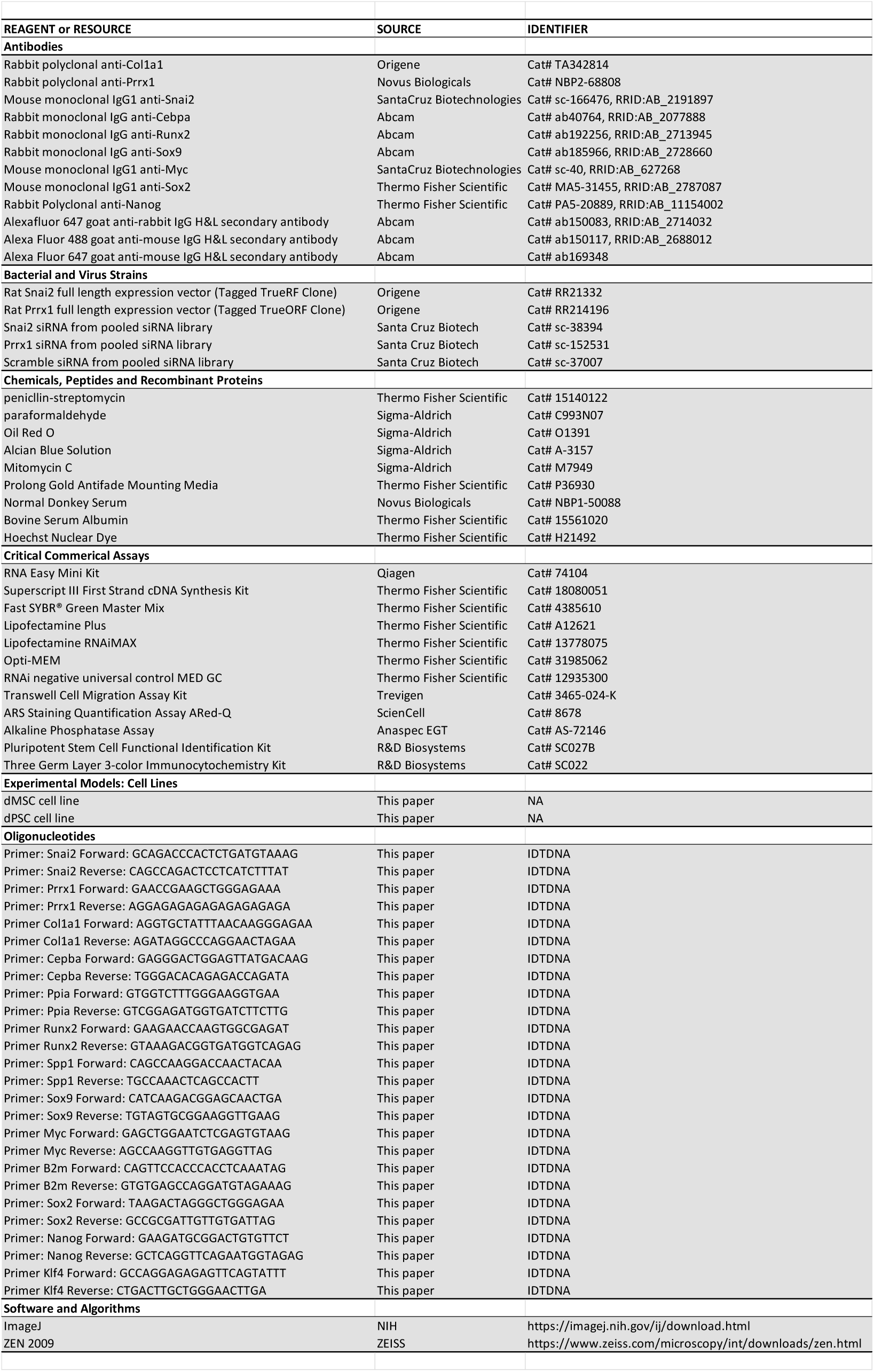

## References

Acharya, P.S., Majumdar, S., Jacob, M., Hayden, J., Mrass, P., Weninger, W., Assoian, R.K., and Pure, E. (2008). Fibroblast migration is mediated by CD44-dependent TGF beta activation. J Cell Sci 121, 1393–1402.

Botchan, M., Topp, W., and Sambrook, J. (1976). The arrangement of simian virus 40 sequences in the DNA of transformed cells. Cell 9, 269–287.

Bulla, G.A., Luong, Q., Shrestha, S., Reeb, S., and Hickman, S. (2010). Genome-wide analysis of hepatic gene silencing in mammalian cell hybrids. Genomics 96, 323–332.

Cacchiarelli, D., Trapnell, C., Ziller, M.J., Soumillon, M., Cesana, M., Karnik, R., Donaghey, J., Smith, Z.D., Ratanasirintrawoot, S., Zhang, X., et al. (2015). Integrative Analyses of Human Reprogramming Reveal Dynamic Nature of Induced Pluripotency. Cell 162, 412–424.

Cao, N., Huang, Y., Zheng, J., Spencer, C.I., Zhang, Y., Fu, J.D., Nie, B., Xie, M., Zhang, M., Wang, H., et al. (2016). Conversion of human fibroblasts into functional cardiomyocytes by small molecules. Science 352, 1216–1220.

Davis, R.L., Weintraub, H., and Lassar, A.B. (1987). Expression of a single transfected cDNA converts fibroblasts to myoblasts. Cell 51, 987–1000.

Ebrahimi, B. (2015). Reprogramming barriers and enhancers: strategies to enhance the efficiency and kinetics of induced pluripotency. Cell Regen (Lond) 4, 10.

Evseenko, D., Zhu, Y., Schenke-Layland, K., Kuo, J., Latour, B., Ge, S., Scholes, J., Dravid, G., Li, X., MacLellan, W.R., et al. (2010). Mapping the first stages of mesoderm commitment during differentiation of human embryonic stem cells. Proc Natl Acad Sci U S A 107, 13742–13747.

Gingold, J.A., Fidalgo, M., Guallar, D., Lau, Z., Sun, Z., Zhou, H., Faiola, F., Huang, X., Lee, D.F., Waghray, A., et al. (2014). A genome-wide RNAi screen identifies opposing functions of Snai1 and Snai2 on the Nanog dependency in reprogramming. Mol Cell 56, 140–152.

Gomes, I.C., Acquarone, M., Maciel Rde, M., Erlich, R.B., and Rehen, S.K. (2010). Analysis of pluripotent stem cells by using cryosections of embryoid bodies. J Vis Exp.

Graf, T. (2011). Historical origins of transdifferentiation and reprogramming. Cell Stem Cell 9, 504–516.

Hikichi, T., Matoba, R., Ikeda, T., Watanabe, A., Yamamoto, T., Yoshitake, S., Tamura-Nakano, M., Kimura, T., Kamon, M., Shimura, M., et al. (2013). Transcription factors interfering with dedifferentiation induce cell type-specific transcriptional profiles. Proc Natl Acad Sci U S A 110, 6412–6417.

Jackson, M., Taylor, A.H., Jones, E.A., and Forrester, L.M. (2010). The culture of mouse embryonic stem cells and formation of embryoid bodies. Methods Mol Biol 633, 1–18.

Liu, K., Yu, C., Xie, M., Li, K., and Ding, S. (2016). Chemical Modulation of Cell Fate in Stem Cell Therapeutics and Regenerative Medicine. Cell Chem Biol 23, 893–916.

Mikkelsen, T.S., Hanna, J., Zhang, X., Ku, M., Wernig, M., Schorderet, P., Bernstein, B.E., Jaenisch, R., Lander, E.S., and Meissner, A. (2008). Dissecting direct reprogramming through integrative genomic analysis. Nature 454, 49–55.

Nashun, B., Hill, P.W., and Hajkova, P. (2015). Reprogramming of cell fate: epigenetic memory and the erasure of memories past. EMBO J 34, 1296–1308.

Polo, J.M., Anderssen, E., Walsh, R.M., Schwarz, B.A., Nefzger, C.M., Lim, S.M., Borkent, M., Apostolou, E., Alaei, S., Cloutier, J., et al. (2012). A molecular roadmap of reprogramming somatic cells into iPS cells. Cell 151, 1617–1632.

Sivaguru, M., Liu, J., and Kochian, L.V. (2013). Targeted expression of SbMATE in the root distal transition zone is responsible for sorghum aluminum resistance. Plant J 76, 297–307.

Srivastava, D., and DeWitt, N. (2016). In Vivo Cellular Reprogramming: The Next Generation. Cell 166, 1386–1396.

Takahashi, K., and Yamanaka, S. (2006). Induction of pluripotent stem cells from mouse embryonic and adult fibroblast cultures by defined factors. Cell 126, 663–676.

Thoma, E.C., Merkl, C., Heckel, T., Haab, R., Knoflach, F., Nowaczyk, C., Flint, N., Jagasia, R., Jensen Zoffmann, S., Truong, H.H., et al. (2014). Chemical conversion of human fibroblasts into functional Schwann cells. Stem Cell Reports 3, 539–547.

Tomaru, Y., Hasegawa, R., Suzuki, T., Sato, T., Kubosaki, A., Suzuki, M., Kawaji, H., Forrest, A.R., Hayashizaki, Y., Consortium, F., et al. (2014). A transient disruption of fibroblastic transcriptional regulatory network facilitates trans-differentiation. Nucleic Acids Res 42, 8905–8913.

Vierbuchen, T., Ostermeier, A., Pang, Z.P., Kokubu, Y., Sudhof, T.C., and Wernig, M. (2010). Direct conversion of fibroblasts to functional neurons by defined factors. Nature 463, 1035–1041.

Xie, X., Fu, Y., and Liu, J. (2017). Chemical reprogramming and transdifferentiation. Curr Opin Genet Dev 46, 104–113.

Xu, J., Du, Y., and Deng, H. (2015). Direct lineage reprogramming: strategies, mechanisms, and applications. Cell Stem Cell 16, 119–134.

Yang, C.S., Lopez, C.G., and Rana, T.M. (2011). Discovery of nonsteroidal anti-inflammatory drug and anticancer drug enhancing reprogramming and induced pluripotent stem cell generation. Stem Cells 29, 1528–1536.

